# Phosphoproteomics of CD2 signaling reveals an AMPK-dependent regulation of lytic granule polarization in cytotoxic T cells

**DOI:** 10.1101/795963

**Authors:** Vanessa Zurli, Tommaso Montecchi, Raphael Heilig, Isabel Poschke, Michael Volkmar, Giuliana Wimmer, Gioia Boncompagni, Gabriele Turacchio, Mario Milco D’Elios, Giuseppe Campoccia, Nicoletta Resta, Rienk Offringa, Roman Fischer, Oreste Acuto, Cosima Tatiana Baldari, Anna Kabanova

## Abstract

The in-depth analysis of costimulatory signaling enhancing the activity of cytotoxic T cells (CTLs) represents a major approach towards immunotherapy development. Here we report that CD2 costimulation plays a critical role in killing by freshly isolated human CTLs, which represent a challenging but valuable study model to gain insight into CTL biology. We show that CD2 triggering critically aids signaling by the T cell receptor in the formation of functional immune synapses by promoting the polarization of lytic granules towards the microtubule-organizing center (MTOC). To gain insight into the underlying elusive mechanism, we explored the CD2 signaling network by phosphoproteomics, which revealed 616 CD2-regulated phosphorylation events in 373 proteins implicated in the regulation of vesicular trafficking, cytoskeleton organization, autophagy and metabolism. Strikingly, signaling by the master metabolic regulator AMP-activated protein kinase (AMPK) represents a functionally critical node of the CD2 network which regulates granule polarization towards the MTOC in CTLs. Granule trafficking is driven by active AMPK enriched on adjacent lysosomes, illustrating a novel signaling cross-talk between vesicular compartments in CTLs. Our results thus establish CD2 signaling as key for regulating cytotoxic killing and granule polarization in freshly isolated CTLs and strengthens the rationale to choose CD2 and AMPK as therapeutic targets to boost CTL activity.

## Introduction

The immune synapse, which is formed between a cytotoxic T cell (CTL) and its target, is characterized by a dramatic intracellular reorganization. In order to kill, CTL must polarize both the microtubule-organizing center (MTOC) and lysosome-like organelles filled with perforin and granzymes, called “lytic granules”, towards the contact site. Lytic granules are transported along microtubules through a dynein-dependent mechanism towards the MTOC, which becomes docked beneath the synapse, to focally release their contents inside the synaptic cleft (1). Notwithstanding the importance of granule polarization in the killing process and the immune surveillance, surprisingly little is known about costimulatory and signaling pathways that regulate granule positioning at the MTOC. The factors implicated comprise the strength of T cell receptor (TCR) stimulation (2), the kinetics of intracellular Ca^2+^ flux (3), and CD103-dependent activation of the phospholipase Cγ1 (PLCγ1) (4). However, the overall knowledge remains fragmentary, and identifying the principal regulators of this process represents an important goal with clear implications for the design of CTL-targeting immunotherapies.

One of the essential tools to gain insight into the biology of CTLs is to study freshly isolated CD8^+^ CD57^+^ T cells that are present at significant percentages in the peripheral blood of healthy human donors (5, 6). The CD8^+^ CD57^+^ effector population is endowed with cytotoxic activity (6, 7), therefore its functioning can be evaluated *in vitro* without prior stimulation or expansion. Importantly, the phenotype and functioning of freshly isolated CTLs appear to reproduce the host phenotype more reliably compared to the *in vitro* expanded CTLs, as exemplified by studies on CTLs from patients and mice with genetic mutations that affect CTL functioning (8). In this context, we previously reported that freshly isolated human CTLs, but not *in vitro* expanded CTLs, form dysfunctional immune synapses with human resting B cells characterized by defective granule polarization(5). Our study highlighted that the functioning of freshly isolated CTLs could be finely regulated during synapse formation by the uncoupling of granule movement from the MTOC translocation towards the synapse, which leads to a dispersion of lytic granules to the CTL periphery with a concurrent loss of cytotoxic killing.

These observations provided us with a robust model to study the signaling pathways controlling the formation of functional lytic synapses between B cells and CTLs and those regulating granule polarization. Here, we addressed both issues by combining two quantitative proteomics workflows. First, we resolved the complexity of the surface proteome of B cells, establishing that CTL costimulation via the surface receptor CD2 interacting with CD58 on B cells was critically important for granule polarization towards the synapse and efficient cytotoxic killing. Then, we explored the outcome of CD2 triggering in CTLs by phosphoproteomics observing that it modulated ∼19% of the CTL phosphoproteome and diverse signaling pathways. Further analysis of the CD2-regulated signaling revealed that the activity of the metabolic regulator AMPK on CTL lysosomes was essential for granule polarization towards the MTOC, defining a new functional role for AMPK signaling in CTLs.

## Results

### Levels of CD58 expression determine the susceptibility of human B cells to lysis by freshly isolated CTLs

Previously we identified several B cell types, including B cells from patients with chronic lymphocytic leukemia (CLL), that were resistant to lysis by freshly isolated CTLs *in vitro* (5). We hypothesized that such resistance was dictated by an altered expression of surface receptor(s) on resistant B cells. To test this hypothesis, we compared cell surface proteomes of resistant and susceptible B cells by using an aminooxy-biotinylation protocol coupled with the streptavidin-based protein pulldown (9). Label-free liquid-chromatography coupled mass spectrometry (LC-MS) analysis allowed us to quantify 1157 putative cell surface proteins, of which 257 were differentially expressed between resistant and susceptible B cells (Figure 1A, Table S1).

**Fig. 1.**
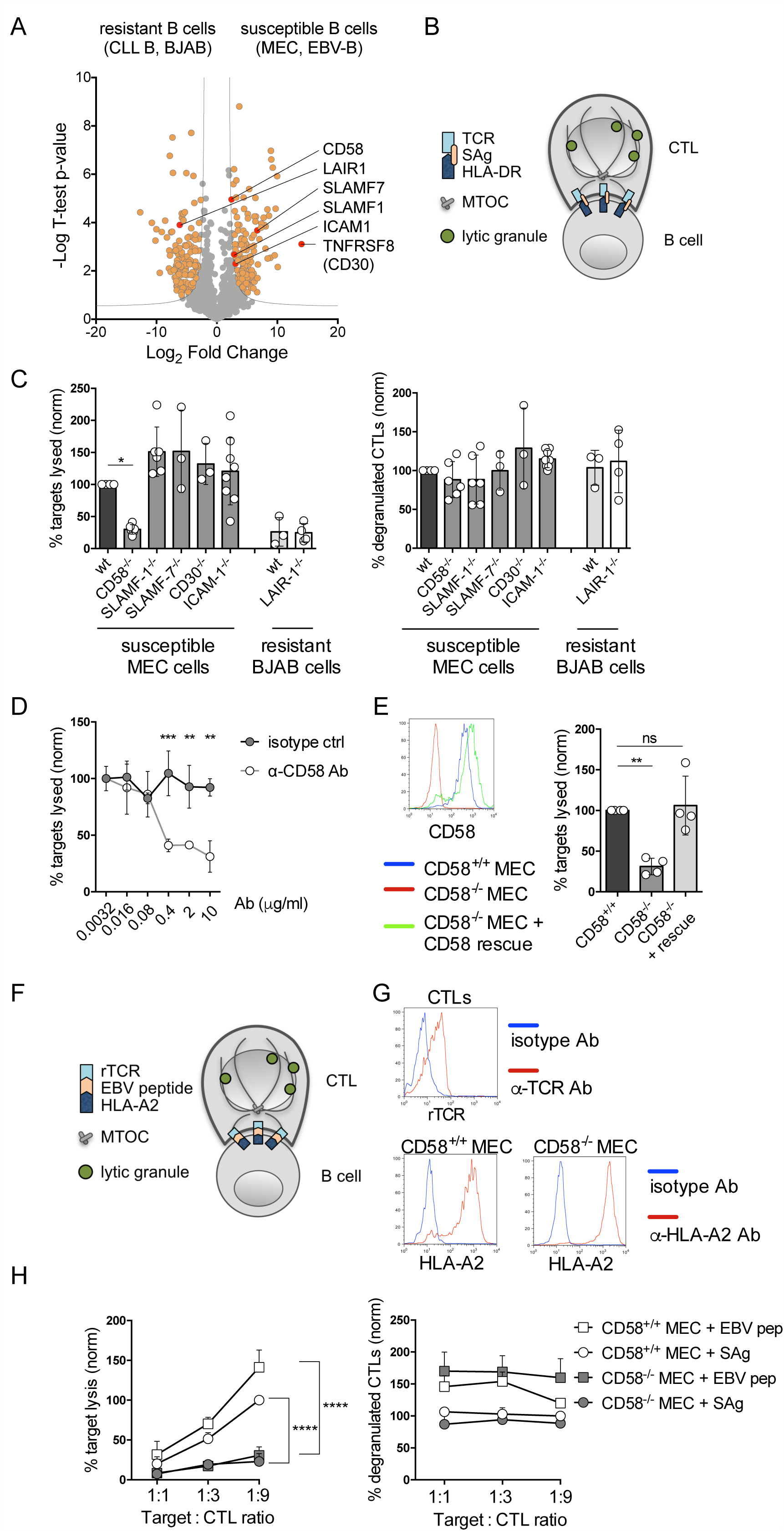
Proteomics discovery of CD58 as the determinant of B cell susceptibility to lysis by freshly isolated CTLs. (A) Surface proteome analysis of B cells, either resistant or susceptible to CTL lysis. Resistant cells were B cells from CLL patients, and a Burkitt lymphoma-derived line BJAB(5). Susceptible cells were the chronic lymphocytic leukemia-derived B cell line MEC, and an Epstein-Barr virus-immortalized B cell line (EBV-B). Volcano plot shows the protein log_2_ fold change (resistant / susceptible B cells) plotted against the -log_10_ *p* value. Proteins with significantly different expression levels are highlighted in orange. (B) Experimental model for the assessment of CTL cytotoxicity against SAg-loaded B cells. (C) Flow cytometry analysis of cytotoxicity and degranulation of CTLs incubated with indicated CRISPR/Cas9-modified MEC and BJAB B cells (n=3-6 experiments). Values for wild-type (wt) MEC cells were set as 100% (killing ranged 10.6-26.3% of targets, degranulation ranged 3.8-8.8% of CTLs). (D) Flow cytometry analysis of CTL killing of MEC cells in the presence of either isotype or CD58-specific blocking antibody (n=3 experiments). Values for the treatment with 0.0032 μg/mL isotype antibody were set as 100% (killing ranged 23.2-41.8% of targets, degranulation ranged 9.6-12.1% of CTLs). (E) Left panel, flow cytometry analysis of CD58^+/+^ and CD58^-/-^ MEC cells, as such or electroporated with CD58 mRNA. Right panel, flow cytometry analysis of cytotoxicity of CTLs incubated with indicated MEC targets (n=4 experiments). Values for CD58^+/+^ MEC cells were set as 100% (killing ranged 9.0-13.3% of targets). For assays (C,D,E) CTLs were incubated with targets at 5:1 ratio. Shown are mean values ± SD; ns, not significant; One- and two-way ANOVA test *p < 0.05, **p < 0.01, ***p < 0.001, ****p < 0.0001. (F) Experimental model for the assessment of killing by freshly isolated CTLs against peptide-loaded B cells. rTCR, recombinant TCR specific for EBV peptide. (G) Flow cytometry analysis of freshly isolated CTLs electroporated with recombinant TCR α (alpha) and β (beta) mRNA, and CD58^+/+^ and CD58^-/-^ MEC cells electroporated with HLA-A2 mRNA. (H) Flow cytometry analysis of cytotoxicity and degranulation of CTLs, electroporated with recombinant EBV peptide-specific TCR mRNA and incubated at indicated ratios with MEC targets, either loaded with SAg mixture or EBV peptide (n=4 experiments). Values for 9:1 ratio of CTLs: SAg-loaded CD58^+/+^ MEC cells were set as 100%. Killing ranged 6.4-28.6% of targets, degranulation ranged 3.1-11.0% of CTLs. Shown are mean values ± SD; two-way ANOVA test ****p < 0.0001.

Among these proteins, we identified five costimulatory receptors highly expressed on susceptible B cells (CD58, CD30, SLAMF-1, SLAMF-7, ICAM-1) and one inhibitory receptor (LAIR1) present at high levels on resistant B cells (Figure 1A). We knocked-out their expression in the CLL-derived B cell line MEC by CRISPR/Cas9-based technology (Figure S1A) to assess whether freshly isolated CD8^+^ CTLs could kill such targets loaded with bacterial superantigens (SAgs) (Figure 1B). We found that only the absence of CD58 had an impact on the killing process, since lysis of CD58^-/-^ MEC cells was decreased by 70.1±7.4 % compared to CD58^+/+^ targets (Figure 1C, left panel; Figure S1B). This effect was recapitulated by a CD58-blocking antibody (Figure 1D). The rescue of CD58 expression on CD58^-/-^ MEC cells led to the restoration of CTL killing (Figure 1E). Notably, even a partial loss of CD58 expression, which occurred as a consequence of heterozygous gene disruption by CRISPR/Cas9, was sufficient to abolish CTL cytotoxicity (Figure S2A), hence indicating that high CD58 levels on target B cells are a critical determinant for cytotoxic killing by freshly isolated CTLs.

To better evaluate the mechanism of CD58^-/-^ MEC cell resistance to lysis, we also analyzed CTL degranulation, which occurs when CTLs interact with antigen-loaded targets (10), by tracking the translocation of lysosome-associated membrane protein 1 (LAMP-1) to the CTL surface during granule release (10). In contrast with the inhibition of cytotoxic killing, the genetic knockout or antibody-mediated blockade of CD58 on MEC cells led only to a moderate decrease in CTL degranulation (20.8±19.6%, Figure 1C, right panel; Figure S2B). This suggested that the levels of interaction between CD58^-/-^ targets and CTLs were largely preserved, however such interaction led to a dysfunctional CTL degranulation associated with the lack of cytotoxic killing.

To confirm our findings also in a physiological setup of CTL-target interaction, we transfected freshly isolated CTLs with a recombinant TCR specific for the EBV peptide GLCTLVAML and assessed their cytotoxicity against peptide-loaded or SAg-loaded CD58^+/+^ and CD58^-/-^ HLA-A2-overexpressing MEC cells (Figure 1F,G). In both settings, the lack of CD58 on targets led to the inhibition of CTL killing and promoted a dysfunctional CTL degranulation (Figure 1H). Overall, our results demonstrate that high levels of CD58 expression on B cells are essential for lysis by freshly isolated CTLs.

### CD2 costimulation promotes the polarization of lytic granules towards the MTOC at the CTL synapse with B cells

Based on our previous findings showing that a defective B cell killing was associated with the loss of granule recruitment to the CTL synapse(5), we hypothesized that the same mechanism might underlie the defect in killing of CD58^-/-^ targets. Confocal imaging analysis revealed that, indeed, the absence of CD58 on target cells led to an impairment in lytic granule polarization to the MTOC docked at the immune synapse, while minimal effects were observed on the polarization of the MTOC itself (Figure 2 A,B). This suggested that costimulation by CD2 regulates granule convergence towards the MTOC. To test this hypothesis, we analysed granule translocation in CTLs incubated with a combination of three anti-CD2 monoclonal antibodies that functionally mimic CD2-CD58 interaction (11, 12) and observed that CTL stimulation via CD2 was able to promote granule polarization towards the MTOC (Figure 2 C,D). Collectively, our data show that CD2 triggering by either a physiological ligand or soluble stimuli is able to strongly promote granule convergence towards the MTOC in freshly isolated CTLs, which in the context of the immune synapse translates into complete polarization of CTLs towards their targets.

**Fig. 2.**
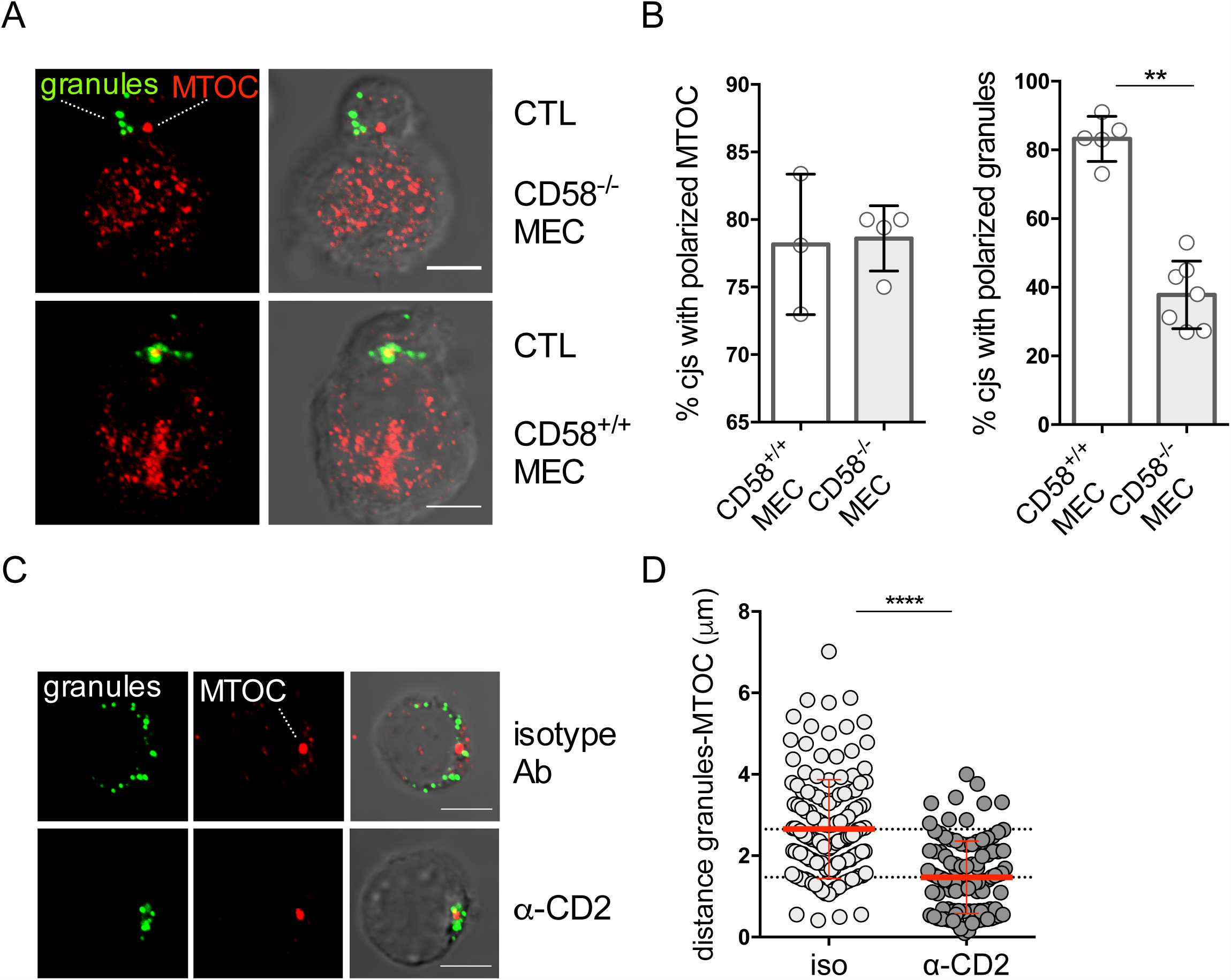
CD2 engagement by CD58 is essential for polarization of lytic granules to the immune synapse of freshly isolated CTLs. (A) Immunofluorescence images of CTLs incubated with CD58^+/+^ and CD58^-/-^ MEC cells (30 min), stained with antibodies against perforin (granules) and gamma-tubulin (MTOC). (B) Left panel: quantitative analysis of MTOC polarization in perforin-positive CTLs incubated with the indicated MEC targets (n=3-5 experiments). Right panel: quantitative analysis of lytic granule polarization in CTLs having the MTOC docked at the synapse with MEC targets. Measurements were taken from 30 conjugates for each condition. (C) Immunofluorescence images of CTLs incubated for 10 min with isotype and CD2-specific (*α*-CD2) antibodies and stained with antibodies against perforin (granules) and gamma-tubulin (MTOC) (n=3). (D) Quantitative analysis of lytic granule polarization in antibody-stimulated CTLs (n=4 experiments). Each dot represents individual granule. Scale bar, 5 μm. Shown are mean values ± SD. Mann-Whitney test **p < 0.01, ****p < 0.0001.

### Phosphoproteomic analysis of CD2 signaling reveals a functionally diversified signaling network of CTL costimulation

To get insights into the signaling pathways of the CD2 network costimulating the formation of CTL synapse and, in particular, those implicated in the regulation of granule polarization, we performed quantitative label-free proteomic analysis of cytoplasmic phosphorylation events in CD2-stimulated CTLs isolated from three individual donors. Analysis of merged samples labeled by tandem mass tags and phosphopeptide enrichment followed by LC-MS revealed a total of 2920 phoshopeptides and 3470 phosphorylation events in 1373 proteins with highly reproducible profile among donors (Table S2, Figure S3). We identified 549 phoshopeptides and 616 phosphorylation events in 373 proteins that were found at significantly different levels in CTLs stimulated with anti-CD2 or isotype control antibody (Figure 3; Table S3).

**Fig. 3.**
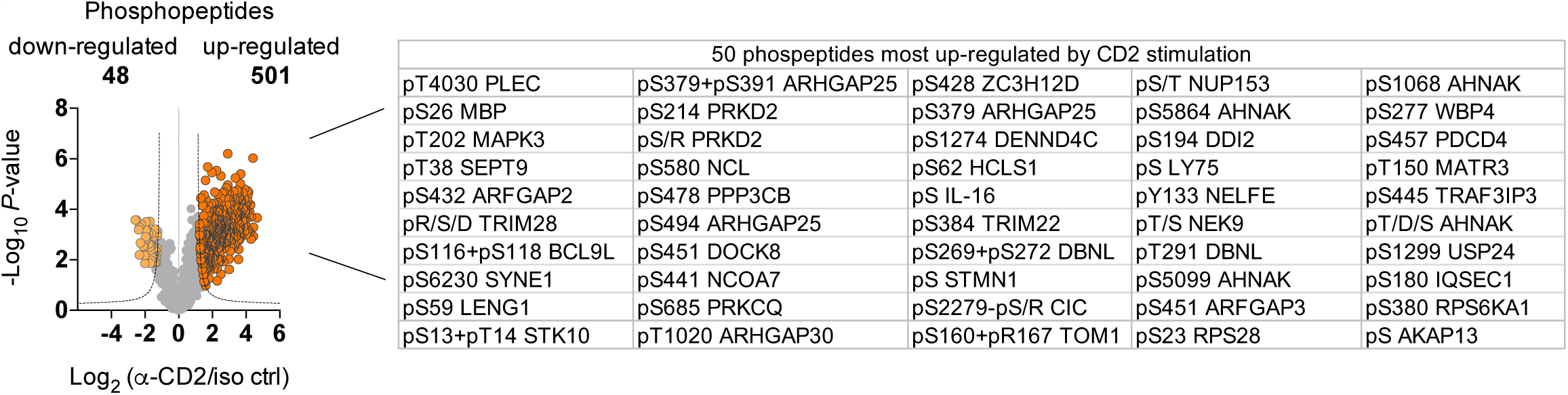
The CD2-regulated phosphoproteome in freshly isolated human CTLs. Volcano plot showes the phosphopeptide log2 fold change (CD2-stimulated / isotype-stimulated) plotted against the -log10 *p* value highlighting significantly regulated phosphopeptides (light and dark orange, n=3). The 50 phosphosites most increased by CD2 stimulation, according to the fold-change in CD2-stimulated vs isotype-stimulated CTLs, are displayed alongside.

The functional diversity of CD2-regulated phosphoproteins was assessed by analyzing the phosphoproteomics data with the Cytoscape software, which allows to create phospho-protein networks based on the interaction data extracted from the STRING database (13). Our analysis revealed that CD2-regulated phosphoproteins formed a complex network of signaling events endowed with several interesting features (Figure S4). First, CD2 regulated the phosphorylation status of a number of phosphoproteins that were predicted to be CD2 interactors (Figure 4A), including several experimentally validated CD2 interactors like CD2AP (14), PTPRC/CD45 (15), and SH3KBP1/CIN85 (16, 17).

**Fig. 4.**
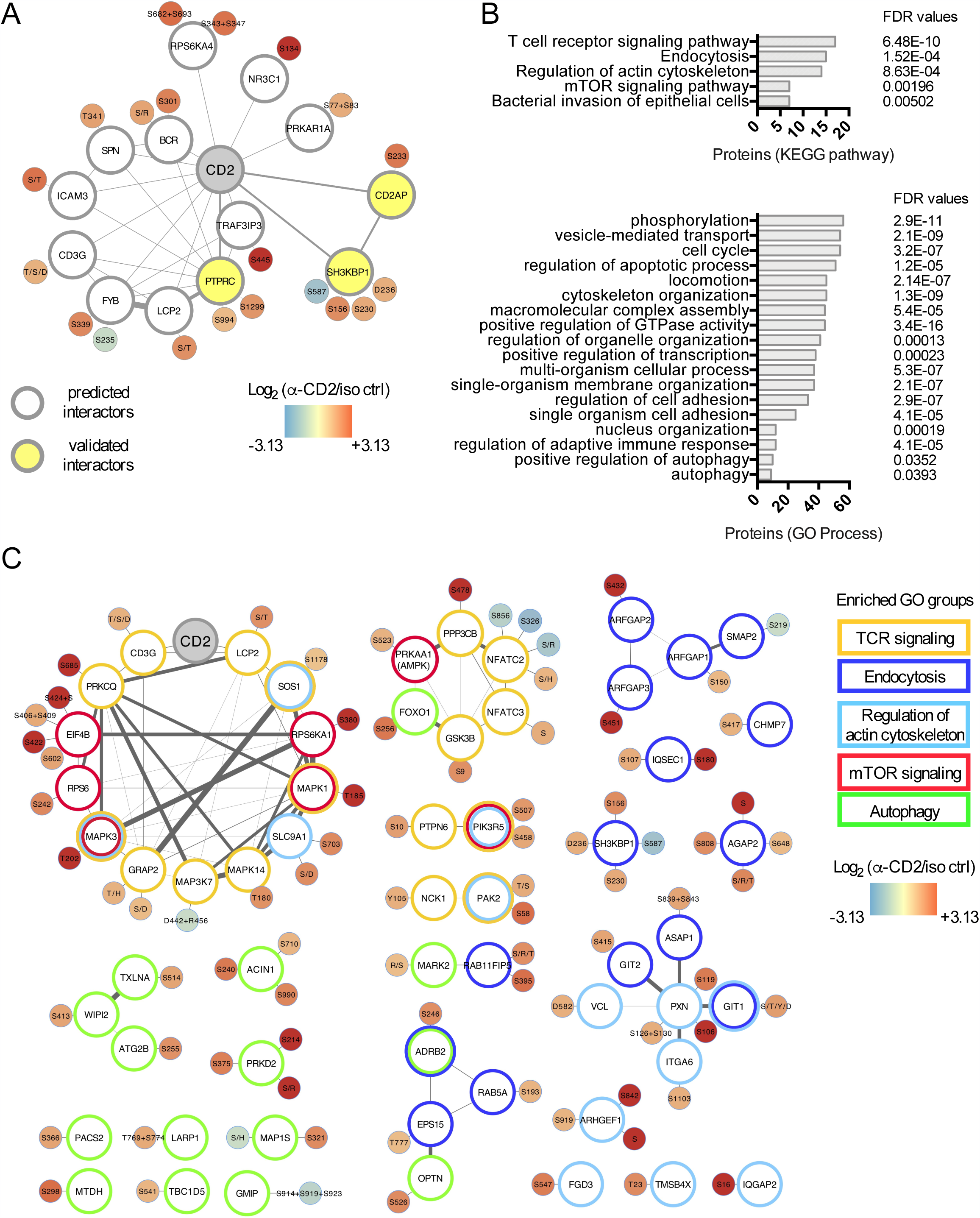
Functional analysis of the CD2-regulated phosphoproteome in freshly isolated human CTLs. (A) A network of validated and predicted CD2-interacting proteins, the phosphorylation status of which is regulated by CD2 stimulation. The network was generated by the STRING application in Cytoscape. (B) Cluster enrichment analysis of CD2-regulated phosphoproteins (Cytoscape application, redundancy cutoff 0.2). FDR, false discovery rate p-values. (C) An integrative network of the CD2-regulated phosphoproteins belonging to the enriched functional groups was generated using STRING application (confidence threshold 0.4) and ClusterMaker application (confidence treshold 0.641) in Cytoscape. For (A) and (C), phosphosite nodes are color-coded based on the phosphopeptide log2 fold change (CD2-stimulated / isotype-stimulated). Edge width is proportional to the confidence of interaction score based on the experimentally validated data from the STRING database.

Further, cluster enrichment analysis of detected phosphoproteins indicated that several functional groups of CD2-regulated phosphoproteins were significantly enriched following CD2 stimulation (Figure 4B, Table S4). Among the enriched signaling pathways, four major pathways included TCR signaling (n=17 proteins), endocytosis (n=15), regulation of actin cytoskeleton (n=14), and mammalian target of rapamycin (mTOR) signaling (n=7). Several relevant functional groups were also enriched, including phosphoproteins implicated in the regulation of cell activation and proliferation (cell cycle n=51, positive regulation of transcription n=38), vesicle-mediated transport group (n=54), locomotion (n=45), cytoskeletal organization (n=45), positive regulation of GTPase activity (n=44), and autophagy (positive regulation of autophagy n=10, autophagy n=9). Altogether, the integrative network of the CD2-regulated phospho-events highlights the ability of CD2 to modulate the phosphorylation status of proteins implicated in a multitude of cellular processes, with particular emphasis on TCR signaling, cell activation, vesicular trafficking and cytoskeleton reorganization, but also metabolic and autophagic processes (Figure 4C).

### The CD2-regulated kinome reveals concomitant activation of the mitogen-activated protein kinase (MAPK) and mTOR pathways

To create a comprehensive map of the CD2-regulated kinome, we merged phosphoproteomics results with the data of immunoblotting and *in silico* prediction of the upstream kinases shaping the CD2-regulated phosphoproteome. First, phosphoproteomics and immunoblotting analyses revealed that CD2 modulated the phosphorylation of 40 kinases (Table S4), including three kinases belonging to the mTOR signaling pathway, namely mTOR (S2448), ribosomal protein S6 kinase beta-1 (p70-S6K) (T389) and AMP-activated kinase (AMPK) (T172) (Figure 5A). Next, we assigned the upstream kinases for CD2-regulated phosphosites by NetworKIN and NetPhorest packages (18). Of 392 phosphosites, for 156 Networkin/Netphorest assigned 41 kinase groups and 82 individual kinases (Table S5). Among those, most phosphorylation events were predicted to be mediated by the PKC group (13.9% of phosphosites), the MAPK group (11.7%), the Akt group (9.9%), and the JNK group (8.8%). In support of this prediction, PKC delta (S304), PKC theta (S685), MAPK1 (T185) and MAPK3 (T202) were found phosphorylated on the activatory residues in our phosphoproteomics analysis (Table S2). Immunoblotting further revealed that Akt1 (S474) and JNK1 (T183/Y185) were phosphorylated on the respective activatory residues (Figure 5B). With this information in hand, we generated an integrative CD2-regulated kinome map, which highlighted intricated molecular interactions and enrichment for the kinases of the MAPK and mTOR signaling pathways (Figure 5C).

**Fig. 5.**
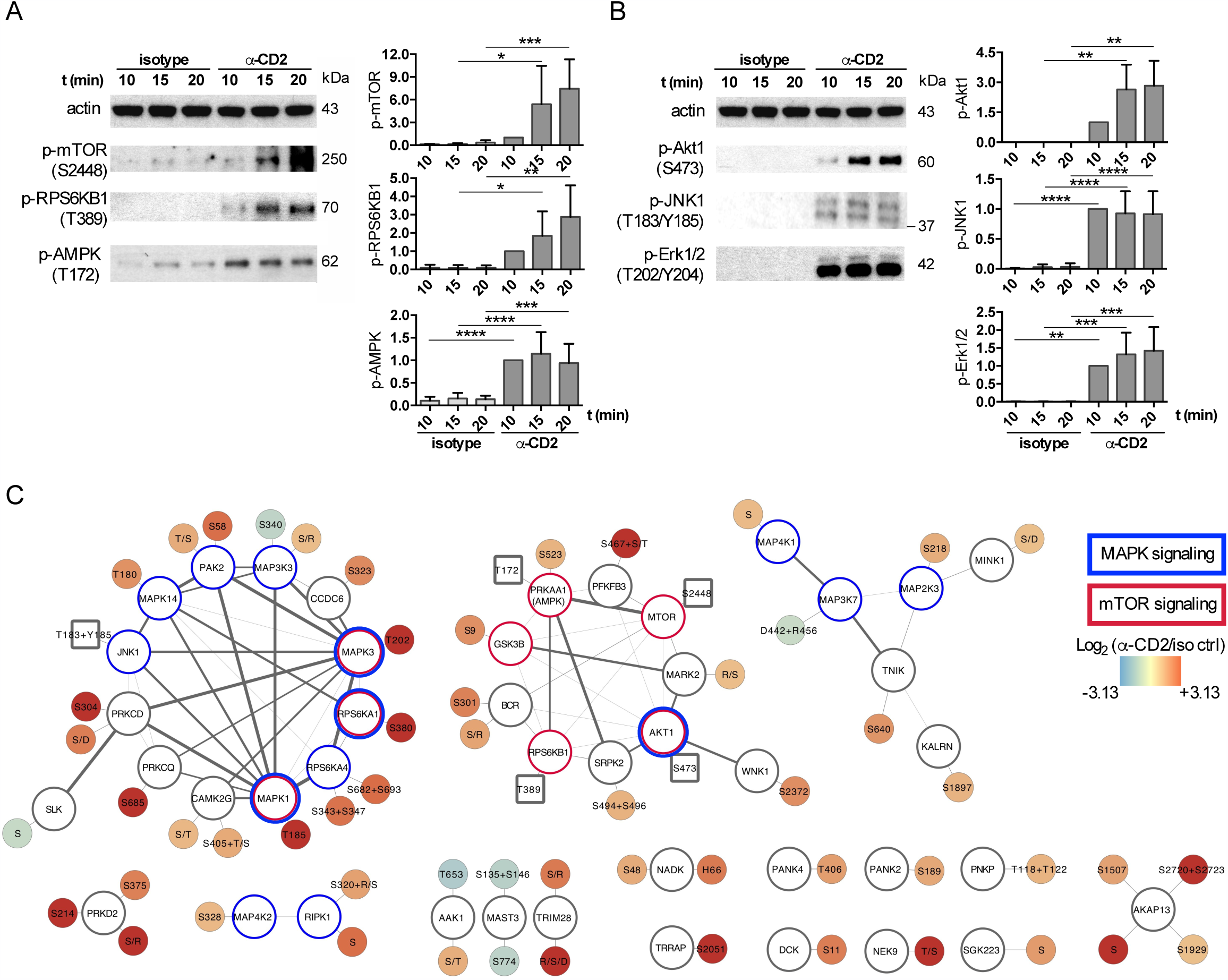
The CD2-regulated kinome in freshly isolated human CTLs. (A, B) Immunoblots of the MTOC signaling pathway (A) and Akt/MPAK signaling pathway (C) in CTLs stimulated either with isotype antibody or CD2-specific antibodies (n=3-4 experiments). Shown are mean values ± SD; two-way ANOVA test *p < 0.01, **p < 0.01, ***p < 0.001, ****p < 0.0001. (C) An integrative network of the CD2-regulated kinases was generated using the STRING application (confidence threshold 0.4) and ClusterMaker (confidence treshold 0.641) in Cytoscape. Round phosphosite nodes refer to the phosphoproteomics data and are color-coded based on the phosphopeptide log2 fold change (CD2-stimulated / isotype-stimulated). Square phosphosite nodes were retrieved from the immunoblotting analysis (Figure S5). Edge width is proportional to the confidence of interaction score based on the experimentally validated data from the STRING database.

### CD2-mediated AMPK activation is essential for granule polarization towards the MTOC

We capitalized on the CD2 phosphoproteomics network to search for signaling events that are responsible for the polarization of lytic granules towards the MTOC. To this end, we concentrated on CD2-regulated lysosome-associated signaling molecules (Table S6). Among these, we found PRKAA1, the alpha 1 catalytic subunit of AMPK which human T cell selectively express (19) and that may have a role in lysosomal trafficking based on a study connecting the metabolic status of the cell with lysosomal positioning (20). We found that CD2 triggering induced PRKAA1 phosphorylation on the activatory residue T172 (Figure 5A) and on the residue S523 (Figure 4C), which was among the top 25% abundant phosphopeptides in CD2-stimulated CTLs (Table S2). Additionally, CD2 modulated the phosphorylation of validated AMPK-specific phosphosites on MFF, GOLGA4, PEA15 as detected by phosphoproteomics (Figure 6A) and on RAPTOR1 and GSK3B as detected by immunoblotting (Figure 6B), confirming that CD2 stimulation promoted an efficient activation of AMPK. Altogether, this motivated our decision to investigate the role of AMPK in granule trafficking.

**Fig. 6.**
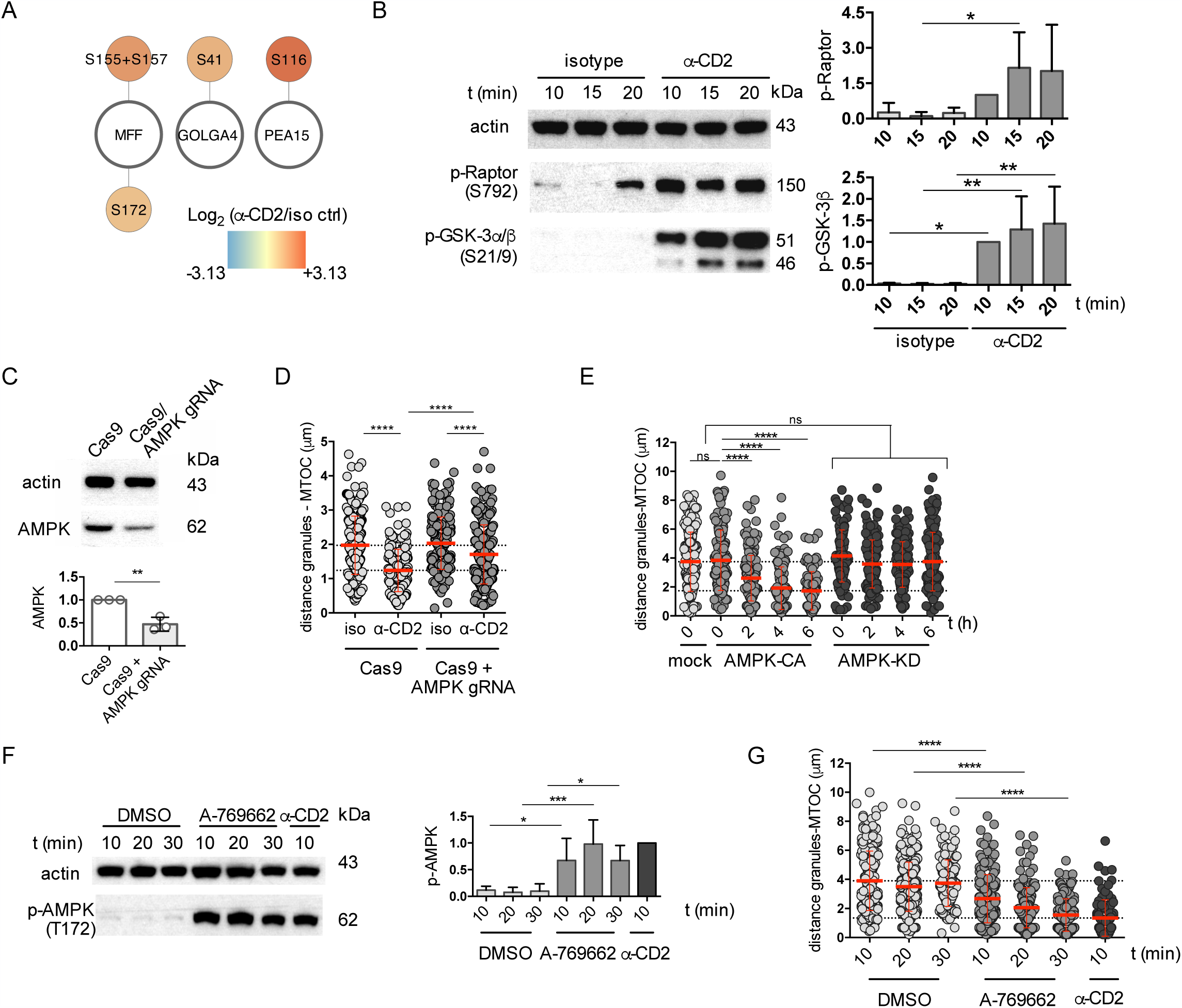
CD2-mediated activation of AMPK is essential for granule polarization towards the MTOC. (A) CD2-regulated AMPK targets identified by phosphoproteomics. Phosphosite nodes are color-coded based on the phosphopeptide log2 fold change (CD2-stimulated / isotype-stimulated). (B) CD2-regulated AMPK targets identified by immunoblotting in antibody-stimulated CTLs (n=3-4 experiments). (C) Immunoblot analysis of CTLs electroporated with RNPs of Cas9 and AMPK-targeting guide RNAs at 48 hours post-electroporation. (D) Quantitative immunofluorescence analysis of granule polarization in Cas9-RNP-electroporated CTLs (n=3 experiments). (E) Quantitative immunofluorescence analysis of granule polarization in CTLs mock-electroporated, or electroporated with mRNA encoding AMPK-CA or AMPK-KD constructs at given time points post-electroporation (n=3 experiments). (F) Immunoblots and quantifications of AMPK activation in CTLs treated with vehicle (DMSO), 300 μM A-769662, or CD2-specific antibodies (n=4 experiments). (G) Quantitative immunofluorescence analysis of granule polarization in CTLs treated with vehicle (DMSO), 300 μM A-769662, or CD2-specific antibodies (n=4 experiments). Shown are mean values ± SD; ns, not significant; two-way ANOVA *p < 0.05, **p < 0.01, ***p < 0.001, ****p<0.0001.

To assess whether AMPK activity was essential for CD2-dependent granule polarization, we used a CRISPR/Cas9 ribonucleoprotein complex-mediated gene knockout to downregulate AMPK expression in freshly isolated CTLs (52.9±15.2% decrease, Figures 6C and S5). This resulted in the impairment of CD2-stimulated granule mobilization towards the MTOC (Figure 6D). These results were confirmed by a small interfering RNA-based approach (Figure S6), collectively showing that the ability of CD2 to induce granule polarization is mediated by AMPK.

To assess whether AMPK activation was sufficient to induce granule polarization in the absence of CD2 triggering, we performed a time course analysis of granule distribution in CTLs transfected with mRNA encoding a constitutively active AMPK mutant (AMPK-CA) or its kinase-dead version (AMPK-KD). This analysis revealed that AMPK-CA construct, but not AMPK-KD construct used as control, promoted granule clustering around the MTOC (Figure 6E). Accordingly, CTL treatment with the AMPK-activating small molecule compound A-769662 induced AMPK activation (Figure 6F) and polarization of lytic granules towards the MTOC (Figure 6G). Altogether, these observations confirm that AMPK activation is the driver of the polarized granule recruitment to the MTOC in freshly isolated CTLs.

### Lytic granule polarization is driven by AMPK localized on lysosomes, but not on lytic granules themselves

Cytoplasmic AMPK accumulates at various subcellular compartments (21). We observed that in freshly isolated CTLs 49.0±18.3% of the intracellular AMPK pool detected by immunofluorescence co-localized with LAMP1-positive lysosomes (Figure 7A,B). To understand whether AMPK activity responsible for granule movement was restricted to the lysosomal compartment, we transfected CTLs with molecular constructs encoding the AMPK inhibitor peptide (AIP) to allow for organelle-specific inhibition of AMPK in lysosomes and in mitochondria, the latter as control (22). This analysis revealed that lysosome-specific, but not mitochondria-specific, inhibition of AMPK completely abolished CD2-induced granule polarization in CTLs (Figure 7C,D), thus suggesting that lytic granule polarization is controlled by lysosome-restricted AMPK activity.

**Fig. 7.**
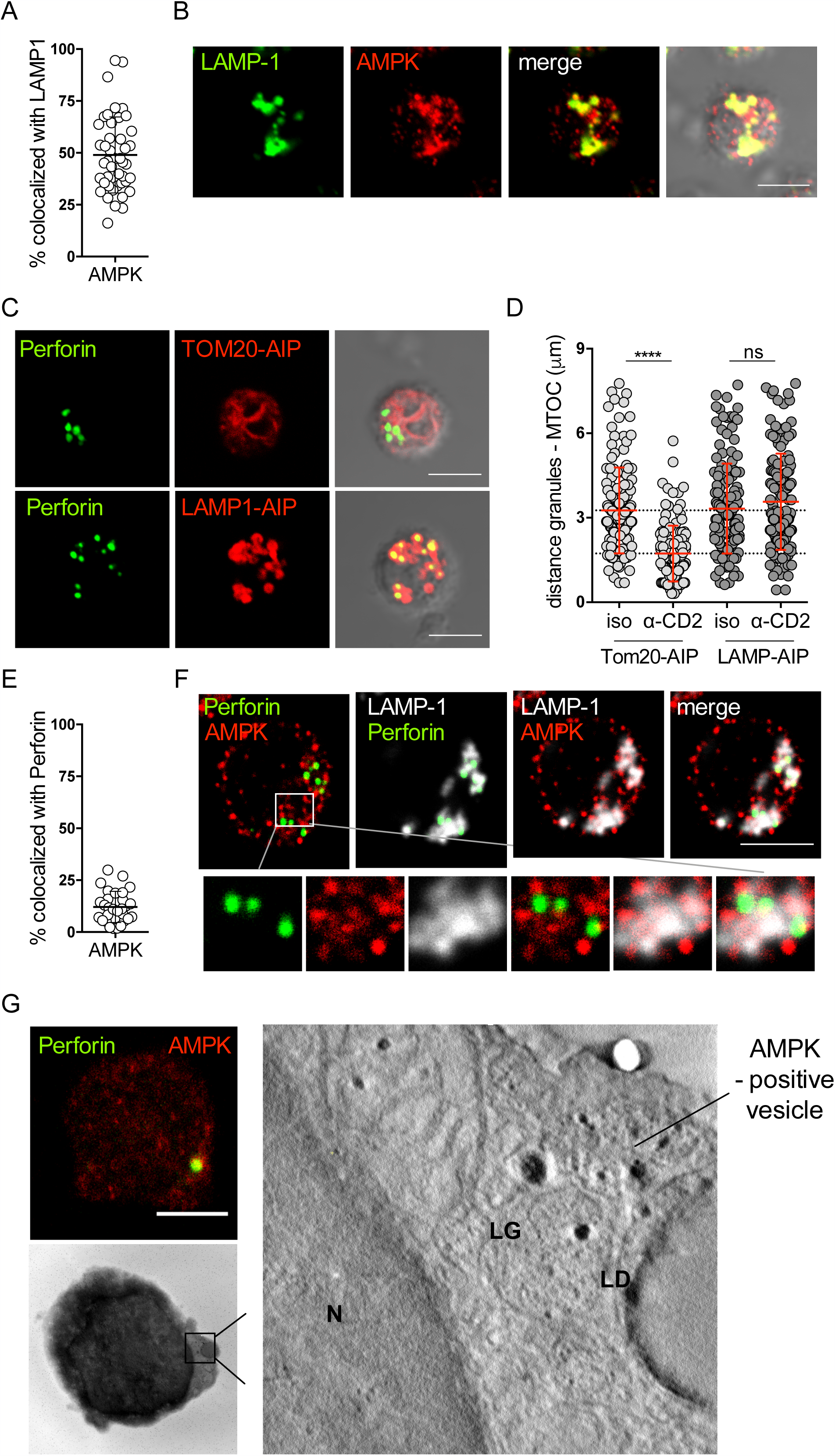
Lytic granule polarization is driven by AMPK localized on CTL lysosomes, but not on lytic granules themselves. (A) Colocalization analysis and (B) immunofluorescence image of LAMP1 and AMPK signals in CTLs. Quantitative measurements were taken for 30 CTLs (n=3 donors). (C) Immunofluorescence images of CTLs electroporated with LAMP1-AIP-mCherry or Tom20-AIP-mCherry constructs and stained with antibodies against perforin (granules). (D) Quantitative analysis of lytic granule polarization in antibody-stimulated mCherry-positive CTLs overexpressing either LAMP1-AIP-mCherry or Tom20-AIP-mCherry (n=3 experiments). (E) Colocalization analysis of immunofluorescence signals in CTLs stained with antibodies against AMPK and perforin (granules) (n=3 experiments). (F) Immunofluorescence image of a CTL stained with antibodies against AMPK, LAMP1 and perforin (granules). (G) CLEM analysis of lytic granules and AMPK-positive compartment in freshly isolated CTLs. N, nucleus; LD, lipid droplet; AMPK, AMPK-positive vesicle; LG, lytic granule. Scale bar, 5 μm. Shown are mean values ± SD; ns, not significant; two-way ANOVA test ****p < 0.0001.

Interestingly, lysosome-associated AMPK did not colocalize with lytic granules themselves, although two compartments were placed in close proximity (Figure 7E,F). These data were confirmed by correlative light and electron microscopy (CLEM) analysis, showing that AMPK-positive vesicles adjacent to lytic granules represented a separate vesicular entity (Figure 7G). Collectively, these observations indicate a functional cross-talk between lytic granules and AMPK-positive lysosomes, whereby local AMPK activity is responsible for the granule polarization process.

## Discussion

In this study, representing an important advance in our understanding of CD2 signaling in human CTLs, we achieve four major goals. First, we provide evidence to the essential role of CD2 in the formation of CTL synapses with B cells, through the costimulation necessary to promote efficient polarization of lytic granules and cytotoxic killing by freshly isolated CTLs. Second, we create a comprehensive map of the CD2 signaling network, which encompasses signaling pathways that regulate cell polarity, vesicular trafficking, cytoskeleton organization, immune and metabolic processes. Third, we identify AMPK as a functionally critical node of the CD2 signaling network responsible for CD2-driven granule polarization in CTLs. Finally, we show that the AMPK pool regulating this process resides on CTL lysosomes, and not on granules themselves, thus illustrating a new example of functional cross-talk between distinct vesicular compartments in CTLs, in addition to the described interaction of lytic granules with Rab27-positive late endosomal compartment required for granule secretion (23).

CD2 has long been known as a potent costimulatory receptor for T cells (24), including CTLs (25, 26), and for another class of cytotoxic lymphocytes, natural killer cells (27, 28). The functional significance of CD2 costimulation applies to pathogen-specific T cell immunity (29) and to autoimmunity (30, 31). In the clinics, in particular, CD2 blockade is highly effective in counteracting autoreactive T cells in diabetes (32), psoriasis (33) and graft rejection (34). Strikingly, CD2 costimulation appears to have an important role also in anti-tumour immunity in B cell malignancies, since the loss of CD58 expression on tumour B cells has been connected with enhanced tumorigenesis in diffuse large B cell lymphoma, non-Hodgkin B cell lymphoma, primary mediastinal large B-cell lymphoma and CLL (35– 38). Our results suggest a molecular mechanism that may underlie these findings, since we show that high CD58 levels on B cells are required for efficient cytotoxic killing by freshly isolated CTLs.

The CD2 receptor has been described as an adhesion and costimulatory receptor on T cells (26, 39, 40), however signaling pathways explaining the potent effect of CD2 costimulation have been elucidated only partially. Known participants in the CD2 signaling network include the TCR CD3ζ subunit (41–43), the kinases Lck (44) and Fyn (45), LAT (46, 47) and WASP proteins (48). Fyn activation by CD2 has been further linked to the activation of the PLCγ1/Vav1/PKC/Dok/FAK/Pyk2/JNK1 axis (45). Here we have characterized the full CD2 signaling network by phosphoproteomics with the principal aim to elucidate signaling molecules aiding the TCR in the formation of functional CTL synapses. First, our analysis revealed the existence of a broad CD2 signaling network in freshly isolated CTLs, which showed a significant overlap with the published signaling phospho-network controlled by the TCR (49) (Table S7), in agreement with the data on TCR transactivation by CD2 previously reported by others (41, 42). Second, we observed that CD2 engagement on CTLs activated several signaling networks highly relevant for CTL synapse assembly, namely, those regulating vesicular trafficking and cytoskeleton organization. Interestingly, these particular functional groups of proteins were among the phosphoproteins that distinguished the phospho-network of CD2 from the published signaling phospho-proteomes of the TCR and CD28 (49, 50) (Tables S7), thus suggesting that CD2 signaling may play a specific role in organizing these processes in CTLs. Also, we found that CD2 regulated the phosphorylation of several proteins previously implicated in the polarization events occurring at the synapse of CTLs and NK cells, including paxillin (S106, S119, S126, S130) (51), Dock8 (S451) (52), and PKC delta (S304) (53) (Table S3).

We were particularly intrigued by our finding that AMPK represents a principal signaling node of the CD2 network responsible for the regulation of granule convergence towards the MTOC in freshly isolated CTLs. Previously, studies performed on *in vitro* expanded CTLs showed that granule polarization at the synapse requires high-affinity TCR-peptide-MHC interaction and rapid kinetics of intracellular Ca^2+^ flux (2, 3), without an apparent need for CD2 costimulation. Our work establishes that freshly isolated CTLs, instead, rely on CD2 costimulation for efficient granule polarization at the immune synapse with B cells, even in the presence of high-affinity TCR engagement by SAgs. We propose that the underlying cause for this could be the ability of CD2 to promote an efficient AMPK activation. In line with this, we observed that in freshly isolated CTLs the CD2-driven AMPK activation is higher when compared to the levels achieved by TCR stimulation (Figure S7). Hence, we propose that in freshly isolated CTLs TCR engagement is essential to promote the interaction between the CTL and its target and to drive the docking of the MTOC at the immune synapse, while CD2 costimulation, through the boost of AMPK activation, serves to ensure complete polarization of lytic granules to the MTOC.

At the molecular level, the involvement of AMPK in lytic granule trafficking in CTLs raises two interesting questions related to the regulation of this process. First, our findings illustrate an example of a direct cross-talk between metabolic signaling and regulation of immune synapse formation. AMPK is the metabolic regulator that in CTLs controls the development of immune response to infections and the formation of immunological memory (54, 55). In the context of anti-tumour immunity, the ability of AMPK to promote granule convergence to the CTL MTOC makes it plausible to propose that AMPK-activating drugs may facilitate the formation of functional CTL synapses whenever granule trafficking is affected by tumour immune evasion. In support of this, we observed that the treatment with the AMPK-activating anti-diabetic compound metformin, known for its anti-tumour activity (56), promotes granule polarization in CTLs (Figure S8). The anti-tumor action of metformin was shown to be dependent on intra-tumoural CTLs, in which metformin counteracts the development of an “exhausted” immunophenotype (57, 58). Our results reinforce these findings by suggesting that this mechanism of metformin action may be aided by the metformin-mediated enhancement of CTL polarization at the immune synapse.

Secondly, an important question is the characterization of molecular events allowing AMPK activation to get translated into the polarized movement of granules towards the MTOC. Changes in the metabolic status of the cell has been implicated in the movement of “conventional” lysosomes, since cell starvation causes lysosome polarization towards the MTOC, that is towards the minus ends of microtubules (20). Driven by the hypothesis that lysosomal positioning could be controlled by the starvation-induced activation of AMPK, in our study we now directly prove that the activation of AMPK is the direct cause of the minus-end translocation of lytic granules in CTLs. The underlying molecular mechanism is as yet elusive. We previously reported that granule positioning in CTLs is regulated by the Arf-like GTPase Arl8 which is the known mediator of the plus-end directed transport of lysosomes (5). When Arl8 expression is downregulated, lytic granules collapse at the MTOC (5), which suggests that the constitutive activity of Arl8 opposes to the constant minus-end directed pulling force and thus maintains lytic granules dispersed. In this framework, we hypothesize that granule polarization may be induced by the CD2-dependent inhibition of Arl8 activity. There is strong experimental evidence supporting this notion, since activated AMPK is known to associate with the LAMTOR/Ragulator complex that behaves as a negative regulator of the Arf-like GTPase Arl8 (59, 60). We hence suggest that CD2-dependent AMPK activation promotes granule polarization via the Ragulator-dependent inhibition of Arl8 activity.

Collectively, here we illustrate an essential role for CD2 and AMPK signaling in the orchestration of CTL synapse assembly with B cell targets, implicating this axis in immune evasion in B cell malignancies. Our data further suggest that CD2 signaling may have a broader function in CTLs, since it modulates the signaling events participating in the regulation of cellular metabolism and autophagy which have emerged as critical regulators of CD8 T cell differentiation and memory formation (61–63). It is also noteworthy that CD2 has the ability to activate two antagonistic metabolism-linked signaling pathways. On the one hand, CD2 promotes mTOR/p70-S6K/Akt1 signaling, which activate anabolic processes. On the other hand, CD2 triggering activates AMPK, which is a key regulator of catabolic processes, including autophagy. It was suggested that a concurrent activation of mTOR and AMPK signaling serves to sustain the rapid induction of biosynthetic processes through autophagy-dependent clearance of misfolded proteins (64) and to achieve high rates of protein secretion (65). Our data thus suggest an interesting functional significance for CD2 signaling implying that it may be directly implicated in the shaping of metabolic, autophagic and secretory profiles in CTLs. A better understanding of this regulation should constitute a basis for the rational manipulation of CD2 costimulation in human pathologies, including hematological disorders.

## Acknowledgements

The authors would like to thank Ellis Reinherz (Dana-Farber Harvard Cancer Center) for the generous gift of CD2-specific monoclonal antibodies; Paul Lehner and James Williamson (Cambridge Institute for Medical Research) for technical advice on surface proteomics; Jim Riley (University of Pennsylvania) for HLA-A2 construct; Veronica Zanon (Istituto Clinico Humanitas IRCCS) for technical advice on T cell culturing; Simona Tavarini (GSK Vaccines, Siena, Italy) for cell sorting. Also, we would like to thank Arthur Weiss, Gillian Griffiths, Morgan Huse, Claire Hivroz, and Juan Bonifacino for fruitful discussions, and Andrés Alcover and Arthur Weiss for the critical reading of the manuscript. This work was carried out with the generous support of AIRC TRIDEO 17015 grant to A.K., and of AIRC grant IG 2017-20148 and ITT-Regione Toscana grant to C.T.B. V.Z. is the holder of an AIRC postdoctoral fellowship.

## Author contributions

V.Z. performed research, analyzed data and wrote the manuscript; T.M., G.W., G.B. performed research and analyzed data; R.H. and R.F. performed proteomics experiments; I.P., M.V., R.O. created transgenic TCR constructs; G.B performed CLEM imaging; M.M.D., G.C. and N.R. provided clinical material and reagents; C.T.B. directed the study and wrote the manuscript; A.K. directed the study, planned and performed the experiments and wrote the manuscript.

## Materials and Methods

### Lymphocyte isolation and culture

Collection of peripheral blood samples of healthy donors was approved by the review board and performed after receiving signed informed consent according to institutional guidelines. CTLs were purified by negative selection using RosetteSep cocktails (StemCell) to purity > 90% for bulk CD8^+^ CTLs and cultured in complete RPMI-HEPES, 7.5% iron-enriched HyClone FCS, 2mM L-glutamine and 50 IU/mL penicillin. For the assays with freshly isolated CTLs we chose donors with at least 15% of perforin-positive cells among total CD8^+^ population (average 23.0±12.3%). If necessary, freshly isolated CTLs were frozen immediately upon isolation and then used after 40-48 h of recovery in complete RPMI-HEPES with 0.01 ng/mL IL-15 (Miltenyi).

### B cell cultures

The CLL-derived B-cell line MEC-1 (MEC) (66) and EBV-transformed B cells (EBV-B) were cultured in complete RPMI (RPMI-1640 with 7.5% iron-enriched HyClone FCS (ThermoScientific), 2mM L-glutamine and 50 IU/mL penicillin). Burkitt lymphoma-derived B cell line BJAB was cultured in complete RPMI medium supplemented with 1 mM pyruvate.

### Reagents

Reagents and antibodies used for immunoblotting, flow cymetry and immunofluorescence are detailed in Supplementary materials. Staphylococcal superantigens SEA (AT101), SEB (BT202) and SEE (ET404) were purchased from Toxin Technology; BSA, poly-L-lysine, propidium iodide, saponin were from Sigma-Aldrich; monensin was from Biolegend; A-769662, STO-609, 5Z-7-oxozeaenol, metformin hydrochloride were from Cayman Chemical; proteomics reagents (periodate, aminooxy-biotin, aniline, lauryl maltoside, iodoacetomide) were from Pierce ThermoFisher.

### CRISPR/Cas9-mutagenesis of B cells and CTLs

Guide RNA sequences are listed in Supplementary methods. Guide RNAs for B cell mutagenesis were cloned into pSpCas9(BB)-2A-GFP (PX458) plasmid (a gift from Feng Zhang; Addgene 48138) following the published protocol (67). B cells were nucleofected with gRNA-encoding vectors or control empty vector using a home-made nucleofection buffer V (90 mM Na2HPO4, 90 mM NaH2PO4, 5 mM KCl, 10 mM MgCl2, 10 mM Na succinate, pH 7.2) and program W-003 on Amaxa Nucleofector II system (Lonza). GFP-expressing cells were sorted, subcloned and screened for gene knock-down by flow cytometry staining. For CTL treatment, 5×10^6^ CTLs were resuspended in 100 μL of Amaxa T cell nuclefection buffer (Lonza) and nucleofected by V-024 pulse with Cas9 RNPs prepared by incubating 5 μg Cas9 (IDT) with 3 μg of *in vitro* transcribed guide RNA (68). CTLs were allowed to recover in RPMI-HEPES with 20% FCS for 48 hours and then used for functional assays.

### Flow cytometry assays of cytotoxicity and CTL degranulation

Assays were done as described(5). Briefly, APCs were loaded with SEA/SEB/SEE, EBV BMFL1_(280-288)_ peptide (IBA GmbH), or 1% BSA for non-stimulated controls. For the cytotoxicity assay, APCs (25×10^3^) were incubated with CFSE-loaded CTLs at the indicated ratio in 50 μL complete RPMI-HEPES for 4h at 37°C. Cytotoxicity was calculated as (%CFSE^-^ propidium^+^ dead cells - %CFSE^-^ propidium^+^ dead cells in ctrl sample)*100/(100 - %CFSE^-^ propidium^+^ dead cells in ctrl sample). For the assay of degranulation, APCs were incubated with CFSE-loaded CTLs in 50 μL complete RPMI-HEPES with 1.2 μg/mL LAMP1-specific antibody (H4A3, Biolegend) for 1 h at 37°C; then 25 μL of monensin (1:333) was added for another 3 h. Fixed cells were permeabilized and stained with Alexa647-anti-mouse IgG(H+L) antibody (Invitrogen) diluted in 0.2% saponin-1% FCS in PBS. CTL degranulation level was calculated as (%CFSE^+^LAMP1^+^ cells - %CFSE^+^LAMP1^+^ cells in ctrl sample)*100/(100 - %CFSE^+^LAMP1^+^ cells in ctrl sample).

### Immunofluorescence microscopy

For immune synapse analysis, 50×10^3^ APCs were loaded with SAg mixture, then washed in RPMI-HEPES and incubated with CTLs (150×10^3^) in 20 μL of RPMI-HEPES at 37°C bath for the indicated time. 15 min before the end of conjugation cells were fixed onto slides in PIPES buffer (69) and stained as indicated. For granule polarization analysis, CTLs (150×10^3^) were left to rest in complete RMPI-HEPES for 10 min, then incubated with anti-CD2 antibodies or 300 μM A-769662 in 40 μL of final volume. Where indicated, CTLs were pre-treated with 10 μM STO-609 and/or 300 nM 5Z-7-oxozeaenol in complete RPMI-HEPES for 45-60 min. At the end of stimulation, cells were washed with ice-cold RMPI-HEPES, fixed onto slides and stained as indicated. Confocal microscopy on 0.9 μm-thick sections was performed on a LSM700 (CarlZeiss) or Leica TCS SP5 using a 63x objective and processed with ImageJ. Distance between lytic granules and MTOC was calculated in at least 30 single CTLs per experiment and condition using ImageJ. Colocalization analysis was performed by calculating Manders coefficient with JACoP plugin in ImageJ.

### Immunoblotting

CTLs (1.5×10^6^) were stimulated or pre-treated with compounds as indicated for the immunofluorescence analysis, then lysed in 40 μL of lysis buffer (20 mM Tris pH 8.0, 165 mM NaCl, 5 mM EDTA, 1% NP-40 supplemented with protease inhibitors (Calbiochem), 1mM Na_3_VO_4_ and 25 mM NaF. Equal amount of post-nuclear cell lysates (10-20 μg) were immunoblotted using primary antibodies and quantified on scanned images with ImageJ.

### Guide RNA sequences for CRISPR/Cas9-based mutagenesis

Guide RNAs for CD58 (GAGCATTACAACAGCCATCG), SLAMF-1 (CGATCTCCTAGATAACGTGG), SLAMF-7 (GAGCTGGTCGGTTCCGTTGG), CD30 (CGGGTCGACATTCGCAGACA), ICAM1 (dual combination TTACTGCACACGTCAGCCGC, CGTGATTCTGACGAAGCCAG), LAIR-1 (TACTGAGTCAATGCGGAATC), AMPK (dual combination AAGATCGGCCACTACATTCT, ATTCGGAGCCTTGATGTGGT).

### Preparation of in vitro transcribed guide RNAs

Guide RNAs were prepared as described (68). Briefly, DNA sequences corresponding to guide RNAs were amplified by using PX458 plasmid as template, the universal reverse primer AGCACCGACTCGGTGCCACT and following forward primers:

1 – TTAATACGACTCACTATAGGAAGATCGGCCACTACATTCTgttttagagctagaaatagc

2 – TTAATACGACTCACTATAGGATTCGGAGCCTTGATGTGGTgttttagagctagaaatagc.

### Validation of AMPK-specific guide RNAs on in vitro expanded CTLs

CTLs blasts were generated from freshly isolated CTLs by using CD3/CD28 dynabeads (ThermoScientific). CTLs were cultured in complete RMPI supplemented with 10 ng/nL IL-2 (Miltenyi), glutamine, penicillin/steptamycin, and 10% FCS. 48 hours after the beginning of expansion, CTLs were depleted of beads, washed once with PBS, and resuspended in buffer 1M (100 μL for 2×10^6^ CTLs). In parallel, Cas9 RNP complexes were prepared by incubating 5 μg Cas9 (Integrated DNA Technologies, 1 μL) with 3 μg of each guide RNA. After 20 min of incubation at RT, RNPs were mixed and used to nucleofect CTL blasts by applying V-024 pulse on Amaxa Nucleofector II (Lonza). CTLs were allowed to recover in complete RMPI with 20% FCS and without antibiotics overnight, and then were cultured as before. AMPK knockout was assessed by immunoblotting 72 hours post-nucleofection.

### CTL nucleofection with siRNA

5-7×10^6^ freshly isolated CTLs were nucleofected with 100 ng AMPK-specific siRNA pool (EHU074041 Sigma-Aldrich) or GFP-specific siRNA control (EHUEGFP) in 100 μL of Amaxa T cell nucleofection buffer (Lonza) and left to recover in 10% FCS-RMPI-HEPES for 40 h before assessing protein knockdown.

### mRNA production and B cell/CTL nucleofection

Full-length CD58 was amplified using as template MEC cDNA (forward primer CTAGCTAGCACCATGGTTGCTGGGAGC, reverse primer CCGCTCGAGTCAATTGGAGTTGGTTCTGTC) and cloned into pcDNA3.1(+). Full-length HLA-A2 was synthetized based on the sequence provided by Jim Riley and subcloned into pcDNA3.1(+) between EcoRV and BamHI sites. EBV-reactive recombinant TCRs (rTCRs) were isolated from a T-cell line raised by repetitive stimulation with the HLA-A2-restricted EBV-derived peptide BMLF1280-288 GLCTLVAML. Using high-precision 96-well sorting on an Aria Fusion cell sorter (BD Biosciences), viable CD3^+^CD8^+^ EBV-reactive T cells were placed at one cell per well into 4.45 µL lysis buffer. Buffer preparation, PCR amplification and sequencing of TCR alpha and beta chains was performed as described(70). Multiple recurrent TCR pairs were selected for cloning and functional testing. TRA and TRB VDJ sequences codon-optimized for human expression were synthesized individually (Eurofins) and introduced using a seamless cloning approach into pcDNA3.1-TRAC and pcDNA3.1-TRBC vectors containing the murine alpha and beta constant regions with an additional disulfide bond(71, 72). BMLF1 reactivity of rTCR was confirmed by exposing rTCR-transduced T cells to targets pulsed with BMLF1280-288 or an irrelevant HLA-A2 restricted peptide. AMPK-CA was amplified using as template pCIP-AMPKa1_WT(73) (a gift from Reuben Shaw, Addgene plasmid # 79010) using forward primer CCGGCTAGCACCatggcgacagccga and reverse primer CCGCTCGAGTTAGTAAAGACAGCTGAGAACTTC and cloned into pcDNA3.1(+). AMPK-CN was amplified using as template pCIP-AMPKa1_KD(73) (Addgene plasmid # 79011) using forward primer CCGGCTAGCACCatggcgacagccga and reverse primer CCGCTCGAGTTATTGTGCAAGAATTTTAATTAGA and cloned into pcDNA3.1(+). For the production of LKB1 mRNA, pcDNA3-FLAG-LKB1 and pcDNA3-FLAG-KD LKB1 (a gift from Lewis Cantley, Addgene plasmids # 8590 and 8591) were used.

For overexpression, mRNA was prepared *in vitro* with 1 μg of XbaI-linearized pcDNA3.1-based construct using HiScribe T7 ARCA mRNA Kit with tailing (NEB). MEC cells were nucleofected with 5 μg CD58 or HLA-A2 mRNA using buffer V and program W-003. CTLs were nucleofected with rTCR alpha and beta pair (5 μg each) using home-made buffer 1M (5 mM KCl, 15mM MgCl_2_, 120mM Na_2_HPO_4_/NaH_2_PO_4_ pH7.2, 50mM Manitol) and program V-024. Post-nucleofection cells were maintained in complete RPMI (10% FCS for T cells) without antibiotics. Overexpression of recombinant proteins was verified by flow cytometry by CD58-, HLA-A2, and anti-mouse TCR beta chain-specific antibodies (to detect rTCR).

### CTL nucleofection with LAMP1-AIP/Tom20-AIP plasmids

For overexpression of LAMP1-AIP and Tom20-AIP, CTLs were nucleofected with 2 μg of purified LAMP-mChF-AIP and Tom20-mChF-AIP plasmids (a gift from Takanari Inoue, Addgene plasmids # 61524 and # 61512).

### Proteomics

#### Enrichment of cell surface proteins of B cells

Enrichment was performed as described(9). Briefly, 120-160×10^6^ B cells (MEC, EBV-B, BJAB and B cells from patients with chronic lymphocytic leukemia) were washed twice in ice-cold PBS and incubated on rocking platform in 3-4 mL of freshly prepared biotinylation/oxidation solution (1 mM periodate, 100 uM aminooxy-biotin, 10 mM aniline) for 30 min at +4°C. Then cell suspension was quenched by 1% glycerol. Cell pellets were washed in 5% FCS-PBS and in PBS-CaCl_2_-MgCl_2_, then were lysed in 3-4 mL of 0.5% lauryl maltoside lysis buffer (150 mM NaCl, 10 mM Tris-HCl pH 7.6). Lysates were supplemented with 5 mM iodoacetomide with protease inhibitors (Roche) and left shaking for 30 min at +4°C. Subsequently, lysates were centrifuged at 2800 xg for 5 min and twice at 16000 xg for 10 min to eliminate nuclei, and then left shaking with high affinity streptavidin agarose resin (Pierce ThermoFisher) at +4°C for 2 h. Streptavidin agarose was pelleted at 1000 xg for 1 min, and washed using Snap Cap columns (Pierce ThermoFisher) 20 times with lysis buffer, then 20 times with PBS/SDS buffer (0.5% SDS). Agarose pellets were reduced in PBS/SDS with 100 mM DTT for 20 min at RT in the dark, washed 3 times with UC buffer (6M Urea, 100 mM Tris-HCl pH 8.5), alkylated in 20 mM iodoacetamide for 20 min in the dark, and washed 17 times with UC buffer and 3 times with 50 mM ammonium bicarbonate pH 8.0. Agarose beads were digested by incubation with trypsin (Promega) ON, following reduction (DTT) and alkylation (IAA) of cysteines. Peptides were desalted on revered phase cartridges (SOLA, Thermo) following manufacturer’s instructions.

#### Phosphoproteomics of CD2-stimulated CTLs: cell stimulation

15 × 10^6^ of enriched effector human freshly isolated CTLs from three donors (67.2, 50.5, 45.7% effector CD57^+^ CTLs respectively) were stimulated for 10 min at 37°C in complete RPMI-HEPES medium with 10 μg/ml of either isotype or a mixture of CD2-specific antibodies (T11.1, T11.2 and HIK27 clones, all from Sanquin) at 5 μg/ml each clone. Cells were pelleted and lysed in 250 μL of 1 % NP40 lysis buffer (150 mM NaCl, 5 mM EDTA and 20 mM Tris-HCl pH8.0). 220 μg of proteins from post-nuclear lysates were reduced in 5 mM DTT, alkylated in 20 mM iodoacetamide and precipitated by methanol-chloroform extraction. Protein pellets were resuspended in 100 mM TEAB and digested with 4.4 μg trypsin (Promega) overnight at 37°C.

#### TMT labeling, HILIC fractionation, and IMAC phosphopeptide enrichement

Desalted peptides (SOLA cartridges, Thermo) were labelled with six channels of a TMT10plex (Thermo) as per manufacturer’s instructions, with exception of using 220 μg total protein/0.8mg TMT reagent. Excess TMT was quenched with 40 mM Tris-HCl. The labelled peptides were pooled, desalted on SepPak Plus cartridges (Waters) and dried down in a vacuum centrifuge. Phosphopeptides were enriched using HILIC/IMAC(74). Briefly, peptides were prefractionated on a HILIC column (Amide 80, 4.6 × 250 mm, Tosoh Bioscience LLC). 30 fractions were collected and concatenated into 10 pools. Fractions were incubated each with 30 μL of a 50% slurry of PHOS-Select Iron affinity gel (Sigma) for 30 min at RT. Samples were washed two times with 0.5 ml 250 mM acetic acid/30% ACN followed by ddH20 and elution with 100 μL of 400 mM ammonium hydroxide.

#### Liquid Chromatography-Mass Spectrometry

Peptides of the surface proteome and phosphoproteomic analyses were separated on an EASY spray column ES803 and analysed on a Dionex Ultimate 3000/Orbitrap Fusion Lumos platform (all Thermo Fisher Scientific)(75). For TMT labelled peptides we used the MultiNotch MS3 method (76) for generating spectra for identification and quantitation of phosphopeptides with parameters as deposited in the data acquisition method (PDX PRIDE(77)). Proteomics raw data are available via ProteomeXchange with identifier PXD013840.

#### Surface proteome data processing

We used label-free quantitation as available in Progenesis QI (Waters) for the differential quantitation of surface-enriched proteins. Data was imported into Progenesis QI using default parameters. MS/MS data was searched with Mascot (Matrix Science, v. 2.5) against the UPR human database (retrieved 07-Sept-2016). Mass tolerances were set to 10ppm for precursor and 0.5 Da for fragment masses. Carbamidomethylation (Cys) was set as fixed, and Deamidation (Asn, Gln) and Oxidation (Met) were set as variable modifications. Peptide level FDR was adjusted to 1% and peptides with a score of <20 were discarded.

#### Phosphoproteomics data processing

LC-MS/MS data were analysed in Proteome Discoverer version 2.2. Proteins were identified with Sequest HT against the UPR human database (retrieved 31-Jan-2018). Mass tolerances were set to 10 ppm for precursor and 0.5 Da for fragment masses. TMT10plex (N-term, K) and phosphorylation (STY) were set as dynamic modifications, alkylation (C) as static modification.

Perseus software was used to perform statistical analysis of proteomics data to identify proteins/phosphopeptides present at significantly different levels in compared samples (log2 fold change ≥1.0). The proteins identified as being regulated by CD2 stimulation were subjected to functional analysis using Cytoscape software, which was used to create protein interaction networks using STRING application (confidence threshold set at 0.4) and ClusterMaker application (MCL cluster, granularity 2.7, STRING score-based edge cut-off 0.641). Cytoscape was also used to annotate the identified phosphoproteins with Gene Ontology (GO) and Kyoto Encyclopedia of Gene and Genomes (KEGG) pathway protein classification, and to perform cluster enrichment analysis (redundancy cut-off set to 0.2). NetworKIN/NetPhorest webtools were used to map identified phosphorylation sites with kinase motifs. For compatibility with NetworKIN, the numbering of phosphorylated residues was manually corrected to correspond to the first protein isoform, since Proteome Discoverer reports phosphosite positions for all potential isoforms of the relevant protein. Therefore, in figures and Tables S3/S6, the numbering for the first protein isoform is shown.

### Correlative light electron microscopy

3×10^6^ CTLs were fixed onto polylysine-coated 35 mm imaging dishes (Mattek) and fixed in 4% PFA/0.05% glutaraldehyde for 10 min at RT, quenched with 0.2 M glycine in PBS and stored in PBS. Fixed cells were permeabilized with 0.1% saponin / 0.1% BSA, blocked with 5% BSA and stained with AMPK-specific and perforin-specific antibodies as described. Cell of interest were then imaged using Zeiss LSM700 confocal microscope along with the image of the Mattek grid that was imaged in phase contrast. The cells were then processed for electron microscopy. Briefly, the cells were stained with osmium tetraoxide (1%) and potassium ferrocyanide (1.5%) to label the membranes followed by uranium acetate (0.5%). The cells were then dehydrated in graded series of alcohol and embedded in Epon 812. The marking in the grid was used to locate the cell of interest which was then sectioned and imaged in FEI Tecnai 12 electron microscope.

### Statistics

Calculation of mean and standard deviation (SD) values and the statistical analysis was performed using the GraphPad software (version 6). One-way or two-way ANOVA test with *post-hoc* Dunnett, Sidak or Tukey test was used for experiments where multiple groups were compared. Mann–Whitney test was performed to compare two experimental groups.

### Reagents

Staphylococcal superantigens SEA (AT101), SEB (BT202) and SEE (ET404) were purchased from Toxin Technology; BSA, poly-L-lysine, propidium iodide, saponin were from Sigma-Aldrich; monensin was from Biolegend; A-769662, STO-609, 5Z-7-oxozeaenol, metformin hydrochloride were from Cayman Chemical; proteomics reagents (periodate, aminooxy-biotin, aniline, lauryl maltoside, iodoacetomide) were from Pierce ThermoFisher.

## Antibodies/reagents used for flow cytometry and cell stimulation

| Reagent specificity | Supplier / clone | Live/fixed (permeabilization) | Dilution | Incubation |
| --- | --- | --- | --- | --- |
| CD8 | Biolegend / HIT8a | live | 1:200 | 30 min |
| CD28 | Biolegend / CD28.2 | live | 1:50 | 30 min |
| CD27 | Biolegend / O323 | live | 1:50 | 30 min |
| CD57 | Biolegend / HNK-1 | live | 1:50 | 30 min |
| CD58 | Biolegend / TS2/9 | live | 1:50 |  |
| SLAMF-1 | Biolegend / A12(7D4) | live | 1:100 | 30 min |
| SLAMF-7 | Biolegend / 162.1 | live | 1:50 | 30 min |
| CD30 | Biolegend | live | 1:50 | 30 min |
| ICAM-1 | Biolegend / HCD54 | live | 1:50 | 30 min |
| LAIR-1 | Biolegend / NKTA-255 | live | 1:50 | 30 min |
| HLA-A2 | Biolegend / BB7.2 | live | 1:50 | 30 min |
| Recombinant TCR (mouse TCR beta chain) | Biolegend / H57-597 | live | 1:50 | 30 min |
| Perforin | Biolegend / dG9 | fixed (PBS-0.1% Triton-1% BSA or MeOH) | 1:50 | 30 min |
| CD2 | Sanquin / T11.1, T11.2, HIK27 | stimulation | 10 µg/mL |  |
| CD2 | Laboratory Ellis Reinherz (T11<sub>2</sub>, T11<sub>3</sub>) | stimulation | 1:100 (ascites) |  |
| CD3 | Biolegend / UCHT1 | stimulation | 10 µg/mL |  |

## Antibodies/reagents used for immunofluorescence

| Reagent specificity | Supplier / clone | Permeabilization | Dilution | Incubation |
| --- | --- | --- | --- | --- |
| Perforin | Biolegend / dG9 | 0.1% Triton, 0.2% saponin or methanol | 1:50 | 45 min |
| LAMP1 | Biolegend / H4A3 | 0.1% Triton or methanol | 1:400 | 1 h |
| LAMP1 in degranulation assay | Biolegend / H4A3 | present in culture supernatant | 1:400 | 40-50 min** |
| γ-tubulin (mouse IgG1) | Sigma / GYU-88 | methanol | 1:200 | 1 h |
| γ-tubulin (mouse IgG2a) | Santa Cruz Biotech / C-11 | Methanol | 1:200 | 1 h |
| γ-tubulin (rabbit Ig) | Santa Cruz Biotech / H-183 | Methanol | 1:50 | 1 h |
| AMPK | Cell Signaling / D63G4 | 0.2% saponin | 1:100 | 1 h |
| Anti-mouse IgG(H+L) | Invitrogen | - | 1:500-1:1000 | 30 min |
| Anti-rabbit IgG(H+L) | Invitrogen | - | 1:500 | 30 min |
| Anti-mouse IgG1/IgG2a | Invitrogen | - | 1:1000 | 30 min |

## Antibodies/reagents used for immunoblotting

| Reagent specificity | Supplier / clone or cat.n. | Dilution | Incubation |
| --- | --- | --- | --- |
| Phospho-mTOR | Cell Signaling / 2971 | 1:1000 | ON |
| Phospho-RPS6KB1 | Cell Signaling / 108D2 | 1:1000 | ON |
| Phospho-AMPK | Cell Signaling / 40H9 | 1:1000 | ON |
| Phospho-Akt1 | Cell Signaling / 9271 | 1:1000 | ON |
| Phospho-JNK1 | Cell Signaling / 9251 | 1:1000 | ON |
| Phospho-Erk1/2 | Cell Signaling / 9101 | 1:1000 | 1 h |
| Phospho-Raptor | Cell Signaling / 2083 | 1:1000 | ON |
| Phospho-GSK-3 alpha/beta | Cell Signaling / D17D2 | 1:1000 | ON |
| AMPK | Cell Signaling / D63G4 | 1:1000 | ON |
| LKB1 | Santa Cruz Biotech / E-9 | 1:1000 | ON |
| actin | Millipore / C4 | 1:5000 | 1 h |

