## Supplementary Table legends, Supplementary Figures for "Phosphoproteomics of CD2 signaling reveals an AMPK-dependent regulation of lytic granule polarization in cytotoxic T cells"

**Supplementary Files** (Zurli et al. CD2 signaling in cytotoxic T cells regulates AMPK-dependent lytic granule polarization and killing of B cells)

Supplemental Tables 1-7, legends.
Supplemental Figures 1-8.

**Supplementary Tables**

Table S1
Comparative label-free quantitative proteomics analysis of B cell surface proteome ($log_2$ fold change ≥1.0; false discovery rate (FDR) ≤0.09). Statistical analysis was performed with Perseus software. Related to Figure 1.

Table S2
Quantification of phosphopeptides in human freshly isolated CTLs, related to Figure 3.

Table S3
Quantification of phosphopeptides present at statistically different levels in CD2-stimulated CTLs compared to isotype-stimulated CTLs (log2 fold change ≥1.0; false discovery rate [FDR] ≤0.01). Statistical analysis was performed with Perseus software. Related to Figure 3.

Table S4
Cluster enrichment analysis of proteins containing CD2-regulated phosphopeptides (Fisher's exact test, FDR ≤0.01). Analysis was performed in Cytoscape. The list was filtered to remove redundant terms, redundancy cut-off 0.2. Related to Figure 4.

Table S5
NetworKIN/NetPhorest Predicted Kinases of the CD2-Regulated Phosphosites. Related to Figure 5.

Table S6
CD2-regulated lysosome-associated proteins in human freshly isolated CTLs. The list was generated through the analysis in Cytoscape.

Table S7
Comparison of the CD2-regulated and published TCR- and CD28-regulated phosphoproteomes (ref. Mayya V, et al. Sci Signal 2009; Tian R, et al. *Proc Natl Acad Sci* 2015). Analysis was performed in Cytoscape.

### Supplementary Figures

#### Figure S1

(A) Flow cytometry analysis of indicated surface receptors on CRISPR/Cas9-modified MEC and BJAB cells.
(B) Flow cytometry analysis showing similar levels of HLA-DR (major histocompatibility class II molecules to which SAgs bind) on CD58$^{+/+}$ and CD58$^{-/-}$ MEC cells.

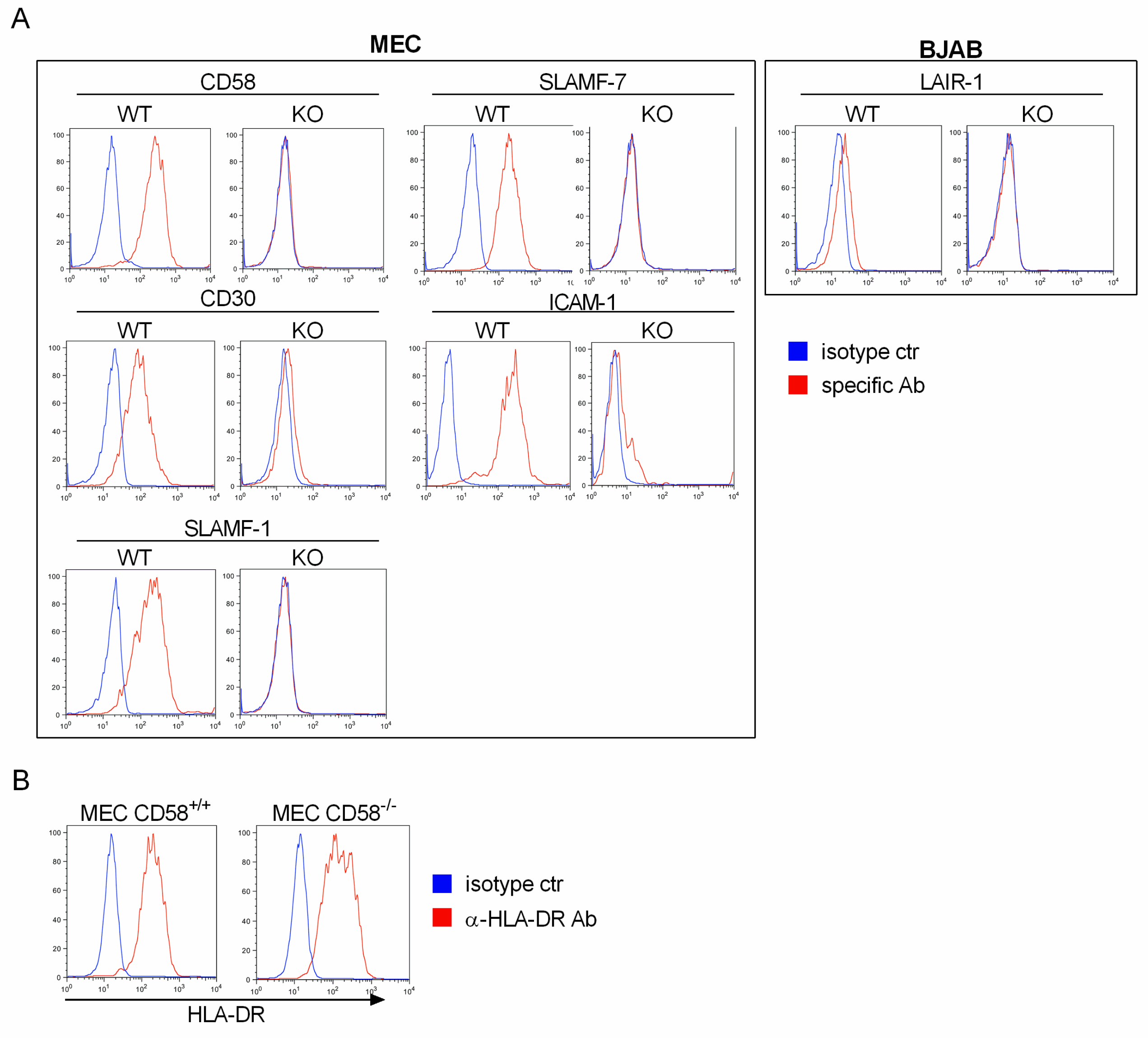

**Figure S2. Influence of CD58 levels on target cells on CTL killing and degranulation**
(A) Left panel: CD58 levels on wild-type MEC cells transfected with empty CRISPR/Cas9-modifying vector (CD58$^{+/+}$), and on CRISPR/Cas9-modified MEC cells with heterozygous (CD58$^{+/-}$) and homozygous (CD58$^{-/-}$) CD58 knockout.
Right panel: Flow cytometry assay of cytotoxic killing with freshly isolated CTLs and indicated SAg-loaded MEC cells.
(B) Analysis of degranulation in CTLs incubated with MEC targets in the presence of either isotype or CD58-specific blocking antibody (n=3 experiments). Normalization was performed setting the values for the treatment with 0.0032 µg/mL isotype antibody as 100% (degranulation ranged 9.6-12.1% of CTLs).

A

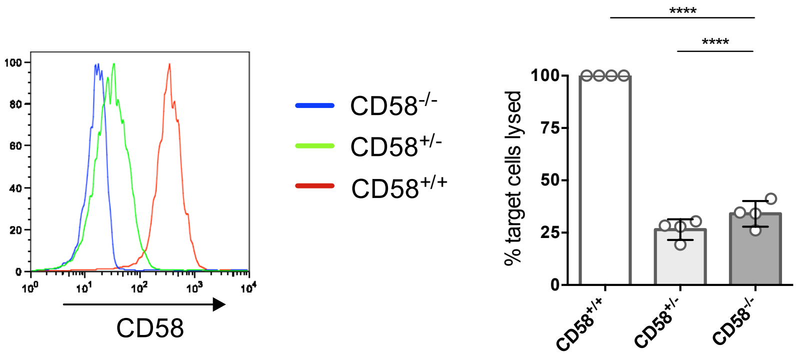


B

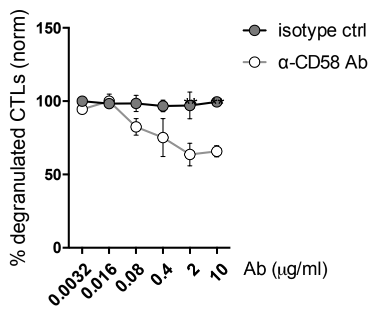

**Figure S3**
Pearson's correlation analysis of CD2-regulated phosphopeptide fold-change quantification among the three donors.

$\alpha$-CD2 Ab

| Donor | 1 | 2 | 3 | |
|---|---|---|---|---|
| | 1 | 0.964 | 0.929 | 1 |
| | | 1 | 0.929 | 2 |
| | | | 1 | 3 |

isotype Ab

| | 1 | 2 | 3 | |
|---|---|---|---|---|
| | 1 | 0.92 | 0.904 | 1 |
| | | 1 | 0.972 | 2 |
| | | | 1 | 3 |

**Figure S4. CD2-regulated phospho-proteome**

An integrative map of 373 CD2-regulated phosphoproteins generated using the STRING (confidence threshold 0.4) and ClusterMaker (confidence treshold 0.641) applications in Cytoscape. Nodes corresponding to kinases are square-shaped.

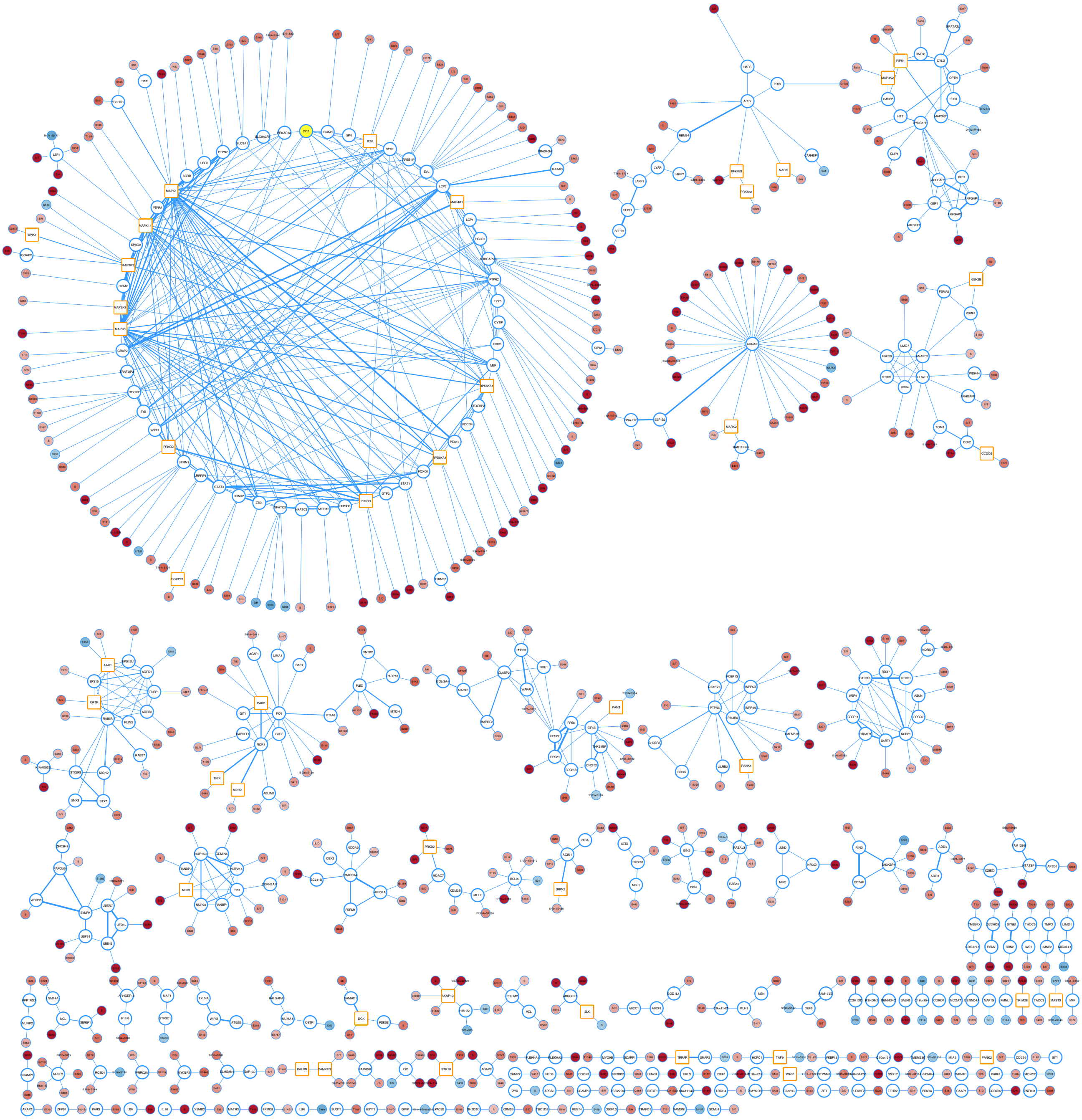

**Figure S5. Validation of AMPK-targeting guide RNAs in in vitro expanded CTLs (CTL blasts)**

Immunoblot analysis of AMPK expression in CRISPR/Cas9-modified CTL blasts at 72 hours post-nucleofection. Cells were nucleofected with Cas9-ribonucleoprotein complexes and several combination of AMPK-targeting guide RNAs. gRNA 1 stands for the combination of GAAGATCGGCCACTACATTC and GGCTGTCGCCATCTTTCTCC guides; gRNA 1+E is the same combination of gRNAs with the addition of a carrier DNA Alt-R Cas9 Electroporation Enhancer (Integrated DNA Technologies); gRNA 2 is the combination of AMPK-targeting gRNAs (AAGATCGGCCACTACATTCT, ATTCGGAGCCTTGATGTGGT) used in the main manuscript.

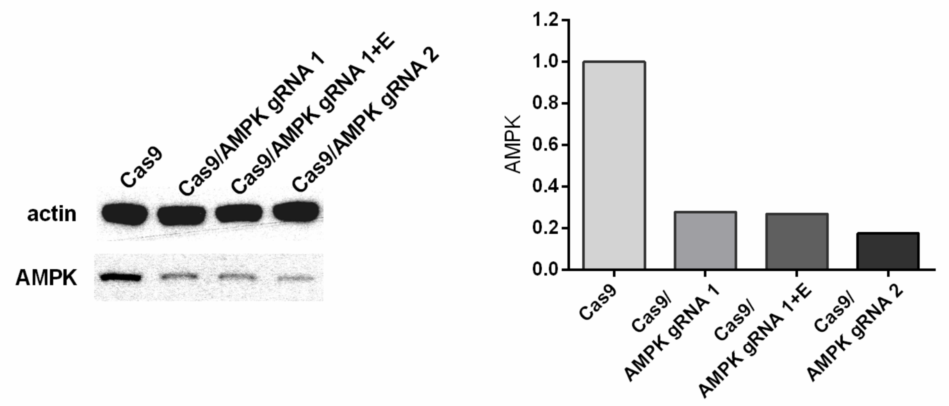

**Figure S6. AMPK knockdown by small interfering RNAs (siRNAs) significantly impaired CD2-dependent granule polarization**

(A) Immunoblot analysis of CTLs electroporated with control siRNA or AMPK-specific siRNA at 40 hours post-electroporation.

(B) Quantitative immunofluorescence analysis of granule polarization in siRNA-electroporated CTLs (n=2 experiments).

Shown are mean values ± SD; ns, not significant; two-way ANOVA test ***p < 0.001, ****p<0.0001.

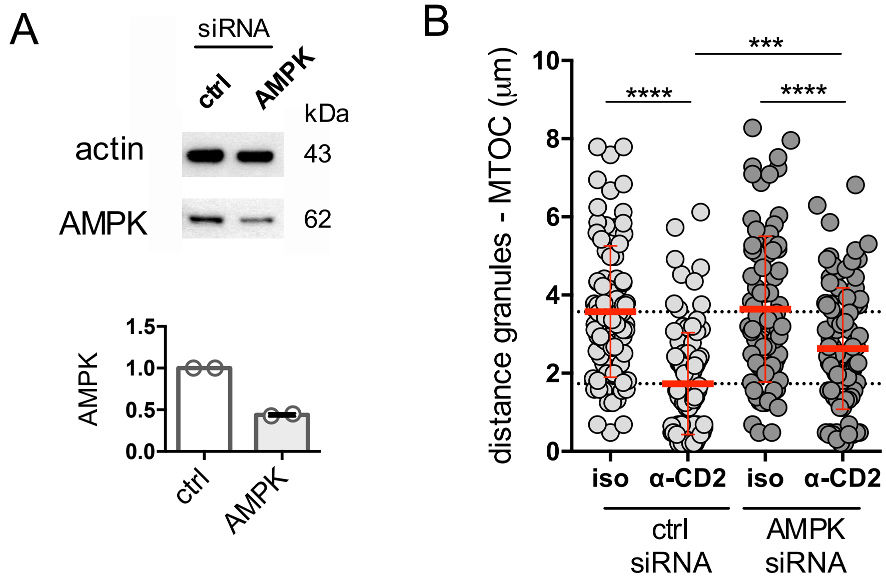

**Figure S7 In freshly isolated CTLs CD2 stimulation triggers enhanced AMPK activation in comparison to the TCR stimulation.**

Freshly isolated CTLs were stimulated with either CD2-activatory antibodies, the TCR-activatory antibody UCHT1, or isotype antibody control (all 10 µg/mL). Left panel: immunoblot analysis of AMPK activation in isotype-, CD2- and CD3-stimulated CTLs (n=4-5). pErk and p-LAT immunoblotting was performed as control blots of the signaling activation. Right panel: immunoblot quantification.

Shown are mean values ± SD; one-way ANOVA test * $p<0.05$, *** $p<0.001$.

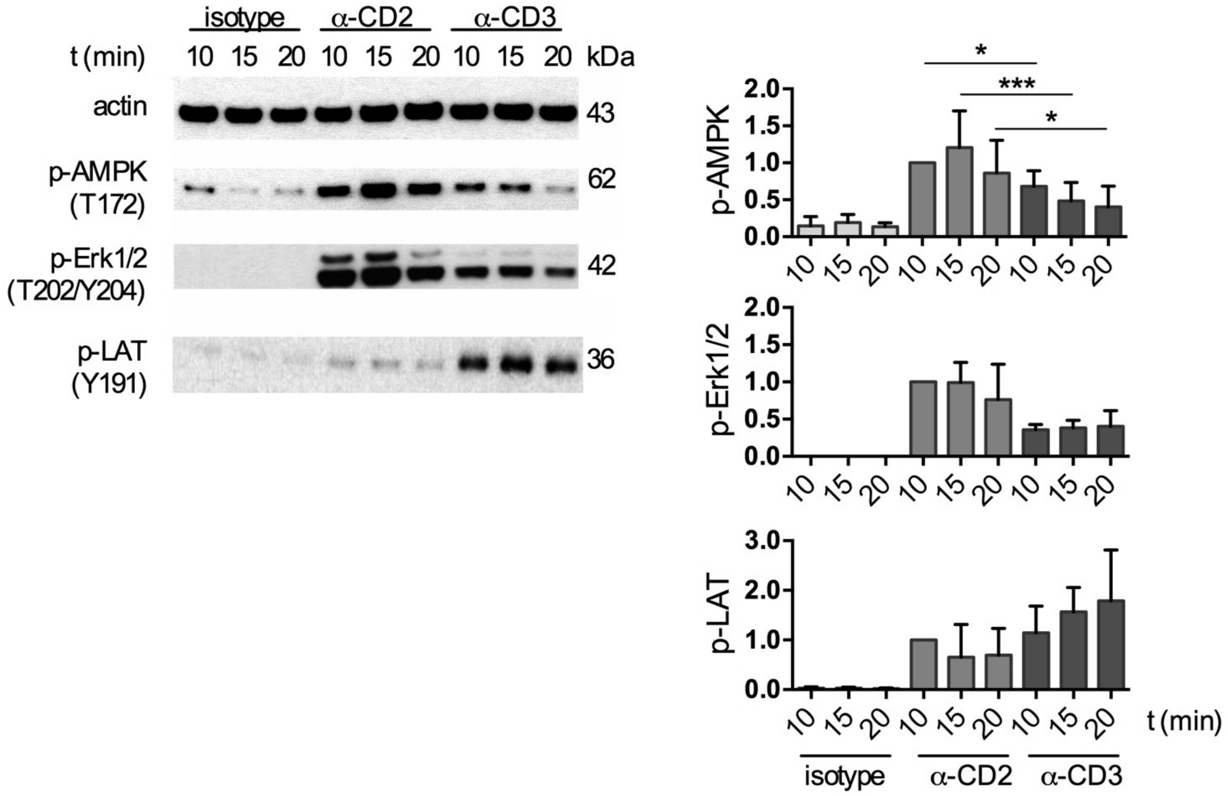

**Figure S8**

Representative quantitative analysis of lytic granule polarization to the MTOC in CTLs treated with vehicle (complete medium), 3 mM metformin, or CD2-specific antibodies (n=2 experiments). Each dot represents individual granule. Shown are mean values ± SD; two-way ANOVA test ****p<0.0001.

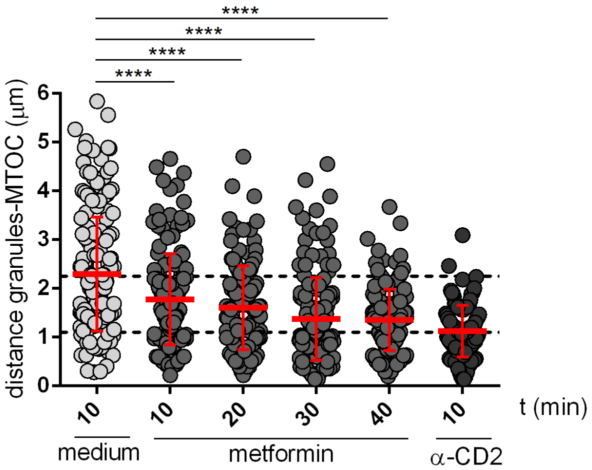
